# Sub-sampling of cues in associative learning

**DOI:** 10.1101/713420

**Authors:** Omar D. Perez, Edgar H. Vogel, Sanjay Narasiwodeyar, Fabian A. Soto

## Abstract

Theories of learning distinguish between elemental and configural stimulus processing depending on whether stimuli are processed independently or as whole configurations. Evidence for elemental processing comes from findings of summation in animals where a compound of two dissimilar stimuli is deemed to be more predictive than each stimulus alone, whereas configural processing is supported by experiments employing similar stimuli in which summation is not found. However, in humans the summation effect is robust and impervious to similarity manipulations. In three experiments in human predictive learning, we show that summation can be obliterated when partially reinforced cues are added to the summands in training and test. This lack of summation only holds when the partially reinforced cues are similar to the reinforced cues (Experiment 1) and seems to depend on participants sampling only the most salient cue in each trial (Experiments 2a and 2b) in a sequential visual search process. Instead of attributing our and other’s instances of lack of summation to the customary idea of configural processing, we offer a formal sub-sampling rule that might be applied to situations in which the stimuli are hard to parse from each other.

## Introduction

The process of generalization is one of the most studied topics in the psychology of learning and decision-making. Of special interest is the problem of *compound* generalization, where an organism needs to predict the consequences of the presence of multiple cues that have independently predicted the same outcome in previous occasions. In this situation, the question at issue is: How much outcome will the organism predict? In the laboratory, the problem is modeled through a summation design, where two cues A and B are separately paired with the same outcome during training and responding to a compound of the two stimuli, AB, is assessed in a final test. A summation effect is obtained when there is more responding to AB than to each of A or B alone (Aydin and Pearce, 1997; Kehoe *et al*., 1994; Soto *et al*., 2009; Thein *et al*., 2008).

The summation effect is readily anticipated by a class of Pavlovian conditioning models called elemental (Rescorla and Wagner, 1972), which assume that subjects represent A and B independently and estimate the outcome of the compound as a linear sum of their individual predictions. This additive generalization strategy makes efficient use of the available evidence under the assumption that the two cues are independent causes of the outcome (Pérez *et al*., 2018). Configural models, by contrast, (Pearce, 1987; Pearce, 1994; Pearce, 2002) assume that subjects process and associate whole configurations with the outcomes that follow; total responding to AB depends on the similarity between the training configurations A and B and the testing configuration AB. As AB shares half of its cues with A or B, organisms should predict an outcome equal to the average of A and B alone(Pearce, 1987), and summation should not be observed.

Experimental evidence on this question in animals has demonstrated a more general role of similarity on summation. Indeed, not only the similarity between the compound AB and each of the components A and B is important, but also that between A and B. The general result is that summation is obtained when A and B are dissimilar, such as when they come from different modalities (e.g., visual and auditory) (Kehoe *et al*., 1994; Thein *et al*., 2008), but not when they are similar, such as when they come from the same modality (for example, visual; Aydin and Pearce, 1995; Aydin and Pearce, 1997). These results have led to a gamut of formal models incorporating flexible representations in which similarity between the elements in a compound makes it more likely to obtain responding to AB closer to the predictions of elemental (i.e., full summation) or configural (i.e., null summation) theories (Harris, 2006; McLaren and Mackintosh, 2002; Melchers *et al*., 2008; Pérez *et al*., 2018; Soto *et al*., 2014b; Thorwart *et al*., 2012; Wagner, 2008).

In contrast with the animal data, evidence for a role of similarity on summation in humans is scarce. In a series of six studies on causal learning, we found no evidence for this hypothesis across several manipulations of similarity between A and B (Pérez *et al*., 2018). Instead, we observed strong and consistent summation, with participants disregarding variables such as color, shape and spatial contiguity when making their predictions. In that paper, we concluded that the simplicity of the visual stimuli allowed participants to easily identify them on the screen, assume them as independent causes of the outcome and sum the individual predictions when presented with the compound AB (Pérez *et al*., 2018). Under such conditions, summation should always follow.

Interestingly, a few studies in humans have shown that factors such as spatial and temporal contiguity might play a role in modulating summation (Glautier, 2002; Glautier *et al*., 2010; Soto *et al*., 2014a). This is consistent with the fact that most of the evidence for configural processing in animals comes from pigeon autoshaping experiments, where animals receive pairings of visual stimuli and food, which results in the animals approaching and pecking the stimuli (Aydin and Pearce, 1995; Aydin and Pearce, 1997). There are two features of this type of experiment that may explain why summation in not obtained. First, the proximity of the pigeon to the screen in which stimuli are displayed limits its ability to sample whole stimulus configurations. Second, the peck’s target is centered on the area dorsalis of the pigeon’s retina (Goodale, 1983; Martinoya *et al*., 1984) which is considered a “second fovea” due to its high density of cells. These two factors suggest that pigeons may be sampling visual information from a circumscribed area (the area predictive of reward), ignoring the rest of the stimulus (Dittrich *et al*., 2010; Soto

*et al*., 2012; Wasserman and Anderson, 1974). Consequently, weak summation may well arise from inefficient sampling of a stimulus compound within a visual search process, rather than from configural processing prompted by strong similarity between components.

A direct implication of this hypothesis is that visual stimuli designed to be more difficult for human participants to be parsed and processed in parallel should promote sequential search and thus weaken the summation effect. To this end, in Experiment 1 we trained complex stimuli where the target stimuli A and B are presented always in compound with other non-target stimuli X and Y, and varied similarity across target and non-target cues. Under a visual search process, the more similar target and non-target cues are the more likely it is that summation will disappear, as participants will only sample one of the stimuli in the compound (Duncan and Humphreys, 1989). By contrast, the more dissimilar target and non-target cues are, the more likely it is that the whole configuration will be processed. In this case, the results should be consistent with the predictions of elemental models in which each element is processed, anticipating a positive summation effect. In Experiments 2a and 2b, we further tested this sub-sampling hypothesis by training target cues A and B with different outcome values and found that the majority of participants reported their predictions for the compound based on the value of only one of the target cues. We present a formal sub-sampling model which can capture both the results of null summation found in our experiments and previous experiments in animals usually interpreted in favor of configural processing of stimuli.

## Results

### Experiment 1

In all experiments in this paper, participants were asked to play the role of an allergist who had to predict the levels of allergy caused by different drugs in a hypothetical patient, Mr. X (see Figure 1). Participants were required to give an assessment of the level of allergy that each drug would cause in a fictitious patient, Mr. X, on a scale of 0 to 35. After clicking with the mouse on a rating scale to make their predictions, participants had to confirm the rating by clicking on a button below the rating scale. Cues were presented 30 times each, in randomized order. Trials were self-paced.

**Figure 1.**
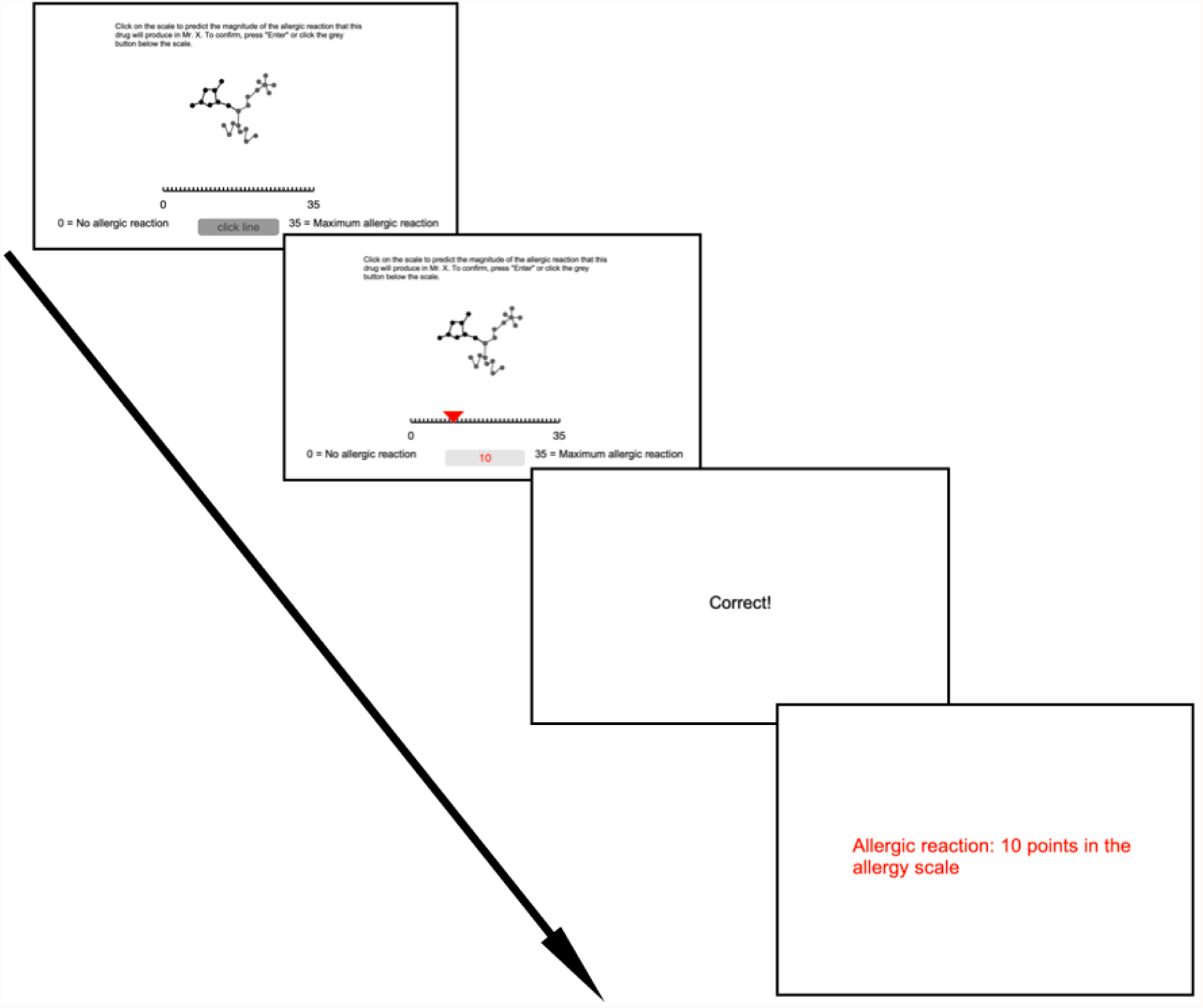
Trial design for all experiments. In each training trial, participants observed one target cue (A,B, C or D) and two non-target cues (X and Y) in the same configuration (AXY, BXY, CXY, DXY) and were asked to rate the level of allergy they thought would be produced by the stimulus on a scale from 0 to 35. After an inter-stimulus interval, feedback was presented (“Correct!” if the prediction was correct, “Incorrect!” if the prediction was incorrect), followed by the message “Allergic reaction: 10 points in the allergy scale” for cues that predicted allergy or “No allergic reaction: 0 points in the allergy scale” for cues that did not predict allergy. During the test phase, the cues were presented in the same way as in training, but now the compound AB, together with one non-target cue (X or Y) was presented in some trials (i.e., ABX, ABY). The predictions were assessed in the same way as in training but no feedback was given to participants during the test phase. Each cue was presented two times during the test.

After confirming the level of allergy expected, participants were presented with a feedback screen with the message “Correct!” if their estimation was correct, or the message “Incorrect!” if the estimation was incorrect (see Figure 1). After this screen, they were presented with the feedback message “10 points of allergy out of a total of 35” for cues that predicted allergy, or “0 points of allergy out of a total of 35”, for cues that did not predict allergy. To encourage learning and attention to the task, each incorrect assessment was followed by a 5-second message (i.e., a time-out); a 1-second message was employed for correct assessments.

The novel addition of this design was the type of training employed. Each training trial included one target cue (i.e., a cue predictive of the outcome; A, B, C or D) and two non-target cues (i.e., cues not predictive of the outcome; X and Y). As can be appreciated in Figure 2, all compounds (AXY, BXY, CXY, DXY, ABX/ABY) consisted of the three cues joined at a central solid circle, with single cues consisting of 8 solid circles each.

**Figure 2.**
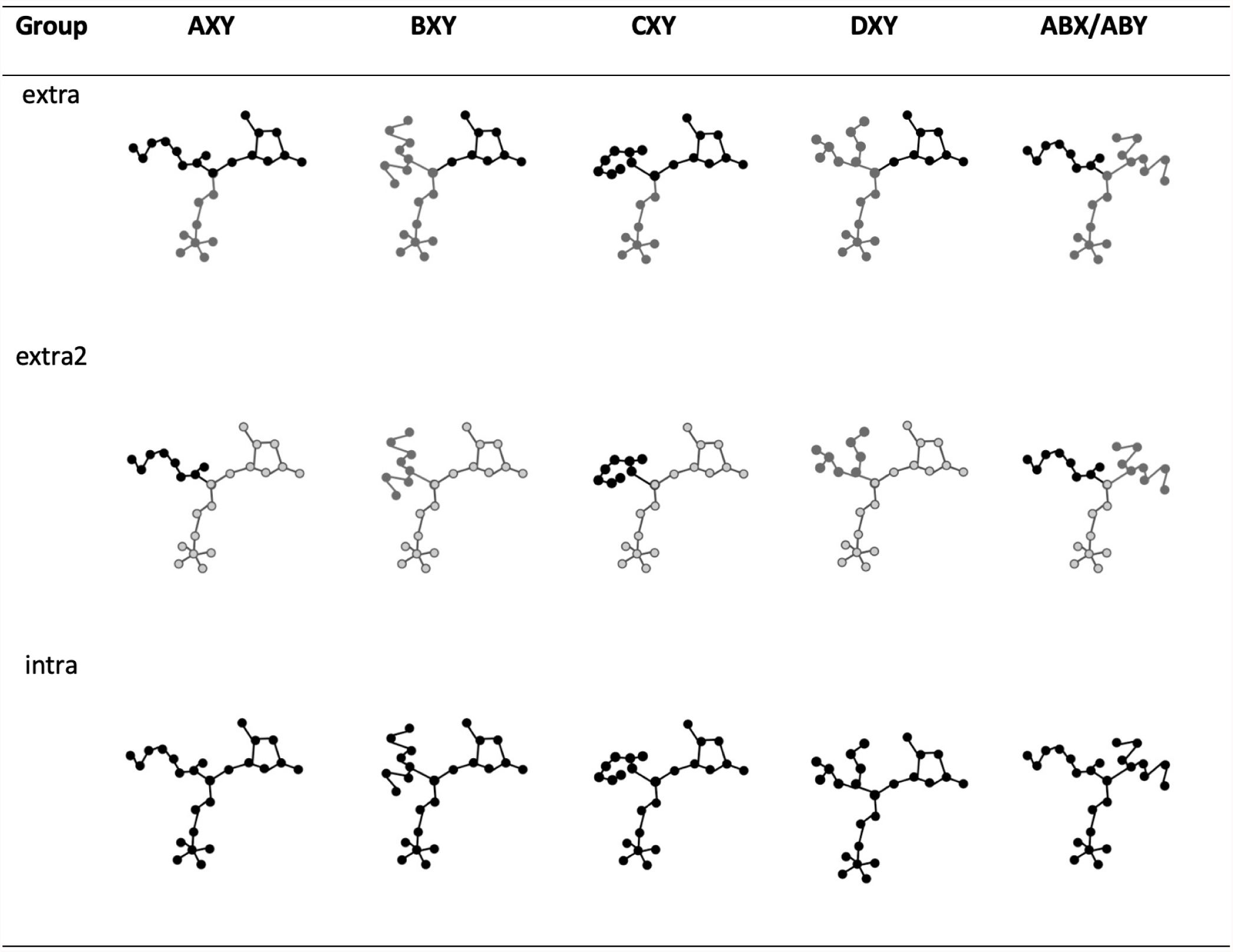
Stimuli used in the experiments reported in this paper. Each stimulus was formed by a central point which branched out into three different components. Target cues A and B were accompanied by two other non-target cues, X and Y. The compound AB was accompanied by one of the non-target stimuli X or Y. All stimuli were presented in three different planar orientations, so that each component was equally likely to appear in one of three positions. Cues A, B, C and D were the target cues. Cues X and Y were contextual or non-target cues that accompanied target cues in each configuration. Stimuli in group *intra* varied only in shape. Stimuli in groups *extra* and *extra2* varied in both shape and color.

Therefore, each compound consisted of 25 circles connected by black lines. In addition, we rotated the compound around its central circle across trials, so that each cue was presented in three possible spatial positions and directions. This ensured that the number of stimuli on each configuration was the same for both training and test phases; the spatial position and direction of the target components did not facilitate their identification.

To test for summation, in some trials of the test phase both A and B were presented in the same configuration, but with only one of the non-target cues (either X or Y, counterbalanced). Cues C and D were included to check which participants correctly learned the contingencies between the cues and their respective outcomes. Because our aim was to investigate generalization of learning, only those participants who correctly identified the contingencies during training (i.e., showed learning) were included in the final analysis. We employed the same criteria as in our previous studies on summation (Pérez *et al*., 2018). Since the cues were designed to be as abstract as possible so as to avoid any prior preference for any of them, the assignment of cues and shapes was kept fixed in all groups.

As can be appreciated in Figure 2, Experiment 1 comprised three groups. Group *intra* included cues that varied only in shape (i.e., intra-dimensional differences). For the two *extra* groups, cues varied both in shape and in color (i.e., extra-dimensional differences). In group *extra*, A and B had different color (black vs gray), but the colors were shared with the non-target cues X and Y (one of them black and the other gray, see top-left stimulus in Figure 2). This means that although A and B can be easily distinguished, they cannot be easily distinguishable from the non-target cues X and Y. Finally, in group *extra2*, A and B had different color (black vs. gray), and they also differed in color with the non-target cues X and Y (light gray with black contours). As a consequence, the similarity between target and non-target cues was lowest in the intra group and highest in the *extra2* group.

Figure 3 sketches the potential of this design to examine the visual search hypothesis. Consider first group *extra* on top of the table. During test with the compounds ABX, ABY in group *extra* one of the target cues shares color with the non-target cue X or Y, which would make the other target cue “pop-out” from the compound (Treisman, 1988). Once the “popped-out” cue is sampled, the search process ends and the participant reports the value of the sampled cue. Under a search hypothesis, therefore, we should not expect summation in this group. The same result is anticipated for group *intra*. For this group, however, the strong similarity between targets and non-targets encourages a serial search strategy. It is expected that participants in group *intra* show a tendency to stop their search and report the rating associated with the first target found, a phenomenon known as *satisfaction of search* in the visual search literature. Such a strategy reduces cognitive and timing costs compared to exhaustive search (Wolfe, 2018). By contrast, the predictions for group *extra2* are in line with those of elemental learning theories. In this case, dissimilarity of all cues results in parallel processing of the two target cues, and reporting of an allergy rating higher than ten (i.e., a summation effect). In the limit, if participants can parse all the cues and process them in parallel without any generalization between them, we should expect a rating of 20 for the compound.

**Figure 3.**
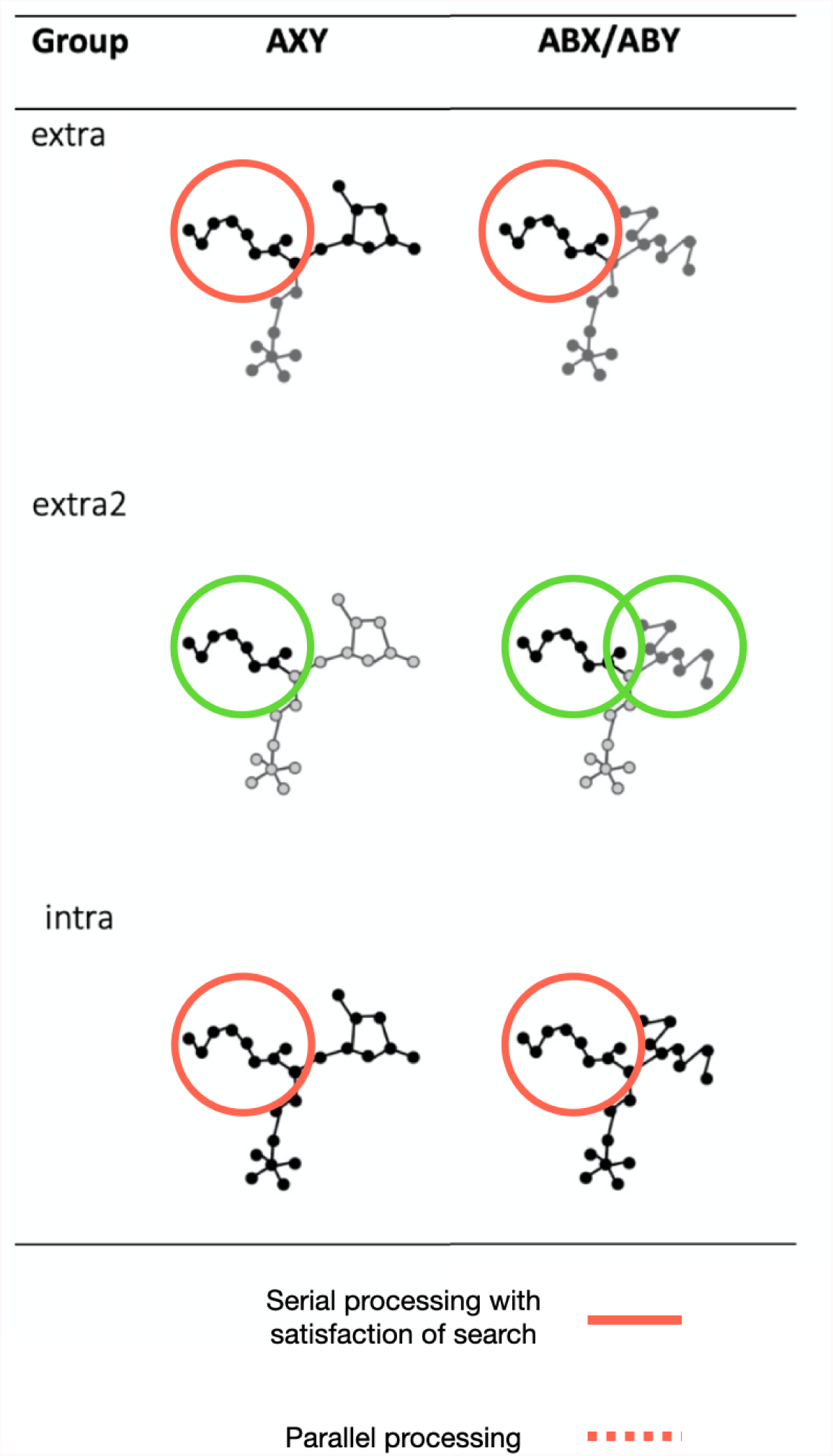
Predictions of a visual search hypothesis for Experiment 1. Under a visual search hypothesis, the difficulty to individuate cues in this design leads participants to perform a visual search for the target cues in the display (Wolfe, 2018) and sub-sample from the whole configuration. Because during training there is always a single target cue in a compound, all groups learn to sample that cue during training (albeit with varied difficulty, depending on similarity between target and non-target cues). Similarity between target and non-target cues fosters serial processing in groups *extra* and *intra*. The result is that participants in these groups would mostly sample a single cue from the testing compounds ABX/ABY and report its associated rating of 10. On the other hand, all cues are processed in parallel in group *extra2*, predicting a rating closer to 20 for the compounds ABX/ABY.

The results are shown in Figure 4, where we present the mean causal ratings to the trained (AXY and BXY) and novel (mean of ABX and ABY) compounds in the three groups of Experiment 1. The figure indicates a summation effect in the form of higher ratings to the novel compounds than to trained compounds in group *extra2*, but no evident summation in groups *extra* and *intra*. A 3 (group) X 3 (compound) mixed design ANOVA confirmed the reliability of these observations in showing a significant group x compound interaction (*F* (2, 55.5) = 4.00, *p* = .02, 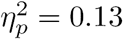, 90%*CI* [.04, .17]). Subsequent simple effects of compound in each group revealed that the mean ratings to ABX/ABY reliably differed from ratings to the trained AXY and to BXY in group *extra2* (*t*(26) = 4.82, *p* < .001, *D* = 0.77, 95%*CI* [2.81, 6.91] and *t*(26) = 4.81, *p* < .001, *D* = 0.77, 95%*CI* [2.83, 6.79], respectively) but did not in groups *extra* (*t*(12) = 0.77, *p* = .46, *BF*_01_ = 2.79 in both cases) and *intra* (*t*(17) = 1.16, *p* = .26, *BF*_01_ = 2.30, and *t*(17) = 1.17, *p* = .26, *BF*_01_ = 2.28, respectively).

**Figure 4.**
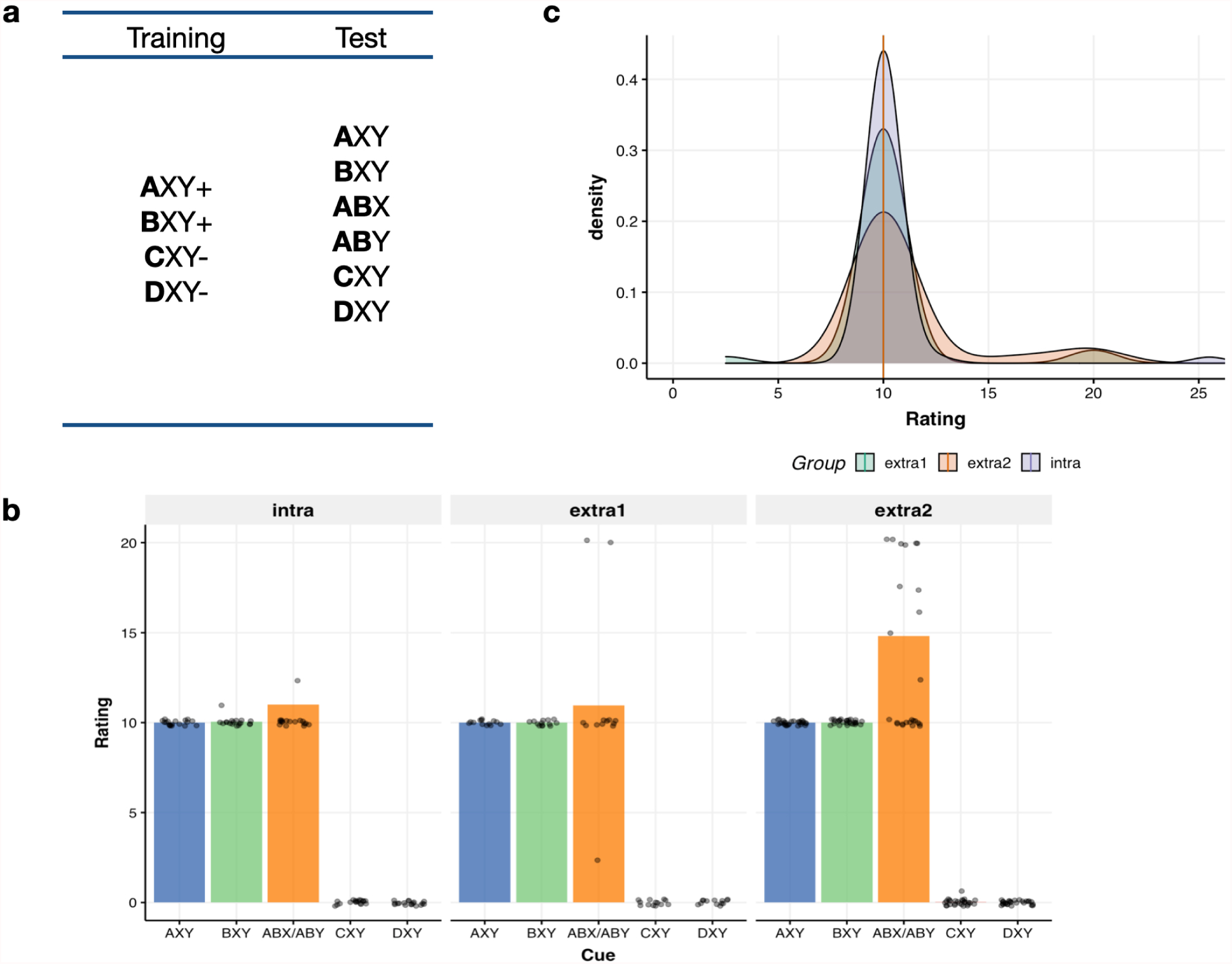
Design and results of Experiment 1. **(a)** Letters denote cues, represented by different chemical shapes that can cause different levels of allergy to a fictitious patient. Each cue is represented by a capital letter. Cues A, B, C and D were the target cues (shown in bold), whereas X and Y were non-target cues. In Experiment 1, all cues were followed (+) or not followed (-) by the same level of allergy (10 points of allergy out of a total of 35). Average ratings given to each cue during the test. Jittered points represent individual ratings. **(c)** Density plots of ratings given to the ABX/ABY cue during the test in the three different groups.

The main effects of compound and group were also reliable (*F* (1, 55.05) = 14.97, *p* < .001, 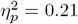, 90%*CI* [.10, .31] and *F* (2, 55) = 3.80, *p* = .03, 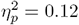, 90%*CI* [.01, .25], respectively), but they are mainly accounted by the summation effect of group *extra2* described above.

To our knowledge, this is the first demonstration of humans being sensitive to manipulations of similarity between target and non-target cues in a predictive learning task. Importantly, we obtained evidence of participants showing no summation in groups *intra* and *extra*, a result that is relatively uncommon in the human literature, as far as we know previously obtained only by Glautier and colleagues (2010). This was achieved by using a design which made it more difficult for participants to distinguish the cues being presented.

We argue that our results are consistent with a serial search process of overt attention which makes participants sample one cue in groups *intra* and *extra* and a parallel search process in group *extra2* where all the cues in the compound are considered in making their predictions during the test with ABX/ABY. We also found that scores for the novel ABX/ABY compound in the different groups followed a bimodal distribution with peaks in 10 and 20, reflecting the fact that participants either find one or both of the target cues to make predictions. This can be appreciated in the distribution of points in Figure 4b, but also in Figure 4c, which shows a density plot of score for the compound ABX/ABY in each group. As can be seen, as similarity between target and non-target cues decreases from *intra* to *extra2*, the proportion of participants processing all cues and consequently showing full (null) summation increases (decreases), with virtually no participants scoring in between these two values, which is what learning theories would predict based on generalization mechanisms. The bimodality of scores was confirmed by Hardigan’s Dip Test: *D* = 0.10, *p* < .001 and a mixture model which yielded a generative model with two Gaussians as the best explanation of the distribution of scores (Gaussian 1: *μ*_1_ = 9.75, *σ* = 2.65; Gaussian 2: *μ*_2_ = 22.10, *σ* = 2.65).

### Experiments 2a and 2b

To further test how sub-sampling of cues can be brought about by visual search, in Experiments 2a and 2b we divided participants in two groups. Group *intra* was a replication of group *intra* of Experiment 1 in which the outcome associated to both AXY and BXY was 10 points of allergy. As in Experiment 1, we expected the majority of participants to rate the compound as producing 10 points of allergy. The critical manipulation was including two different outcome values for AXY and BXY in group *intra2*. In Experiment 2a, we assigned 10 points of allergy for cue AXY and 8 points of allergy for cue BXY; this assignment was reversed in Experiment 2b. To the extent that our stimulus manipulation prompts a search process and limited sampling and processing of a single cue, the majority of participants in group *intra* should rate the compound ABX/ABY as producing either 8 or 10 points of allergy, indicating that they have responded in accord with the value of only one of the two components. More importantly, the value reported by participants in group *intra2* should be the opposite to what participants in group *intra* report. For example, if the features of stimulus A within ABX/ABY are somehow more salient than those of stimulus B, participants in group *intra2* should report 10 points of allergy for ABX/ABY in Experiment 2a for and 8 points in Experiment 2b.

Given that the goal of Experiments 2a and 2b was to test whether participants rate the ABX/ABY compound similarly to either AXY or BXY, we focus our analysis on the distribution of scores and the non-parametric Friedman’s test. Figure 5b presents the barplot and individual scores shown in jittered points for the trained (AXY and BXY) and novel (ABX and ABY) cues in the two groups of Experiment 2a. As expected, most participants rated AXY and BXY in accord with their trained outcomes (10 for AXY in both groups, and 8 and 10 for BXY in groups *intra* and *intra1*, respectively). In group *intra*, median scores were virtually identical for cues AXY, BXY and ABX/ABY with very little variation around 10 points of allergy, which led to no reliable differences in the three distributions (*X*^2^(2) = 0.75, *p* = .16). Differences were apparent in group *intra2* (*X*^2^(2) = 44.69, *p* < .001), where the scores to ABX/ABY (med=8.5) were closer to BXY (med=8) than to AXY (med=10). Dunn-Bonferroni post-hoc tests indicated that the distribution of ABX/ABY differed significantly from both AXY (*p* = .01) and BXY (*p* =< .01) after Bonferroni adjustments.

**Figure 5.**
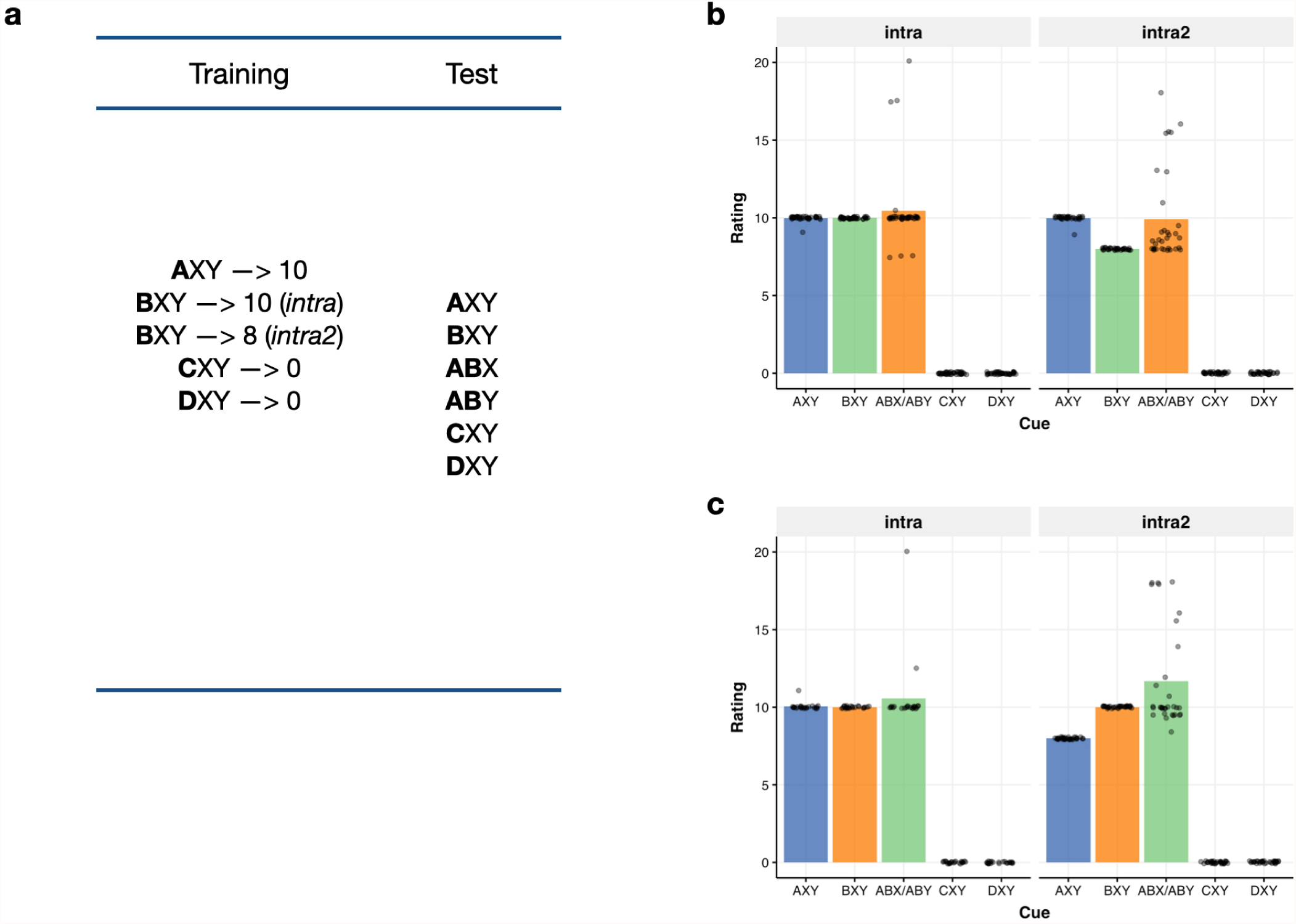
Design and results of Experiments 2a and 2b. **(a)** In Experiments 2a and 2b, cues were followed by different levels of allergy, which are represented by the numbers shown next to each of them. The only difference between Experiments 2a and 2b was the assignment of different levels of allergy outcome to cues AXY and BXY in group *intra2* (8 or 10 points of allergy, counterbalanced across the two experiments). **(b)** Barplots and individual ratings (shown in jittered points) given to each cue during the test in Experiment 2a. **(c)** Barplots and individual ratings (shown in jittered points) given to each cue during the test in Experiment 2b.

The results of Experiment 2b are summarized in Figure 5c. As can be appreciated, the outcome of this experiment is basically the mirror image of Experiment 2a: In the case of group *intra*, we replicated for the third time the absence of any reliable difference among the cues (*X*^2^(2) = 2.46, *p* = .30) which varied in a very small range around 10 points of allergy. In contrast, in group *intra2* the Friedman’s test indicates reliable differences among the cues (*X*^2^(2) = 55.00, *p* < .001), where the ratings to ABX/ABY (med=10) were again very similar to BXY (med=10) and different from AXY (med=8) (*p* = 1.00 and *p* < .001). Thus, ratings to the novel compounds ABX and ABY seem to be determined by the predictive value of B. This change in the distribution of ratings from Experiment 2a to Experiment 2b is not anticipated by any theory of associative learning and is consistent with a visual search approach where participants sample only one of the two target cues to make their predictions of the compound.

### A model of limited cue sampling

Taken together, these results provide evidence for the hypothesis that a visual search process can bring about sub-sampling of cues and therefore weak to no summation in humans when cues are difficult to be parsed and distinguished in a compound.

The purpose of this section is to show how animal results which have traditionally been conceived of as being a consequence of configural processing, may also be explained by sub-sampling of stimuli within a complex visual configuration, such as those employed in pigeon autoshaping where most of the evidence for configural processing, and in particular lack of summation, are found. We present simulations of a formal model based on this idea.

### Learning rule

The model we propose learns according to reinforcement prediction-error (Bush and Mosteller, 1951). The predictive value or *associative strength* of stimulus *i* in trial 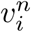, is updated in accord with Equation 1

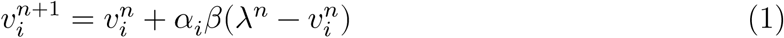

This algorithm assumes that the change in the predictive value of stimulus *i* is determined by the difference between the observed outcome and the current outcome expected from that stimulus.

### Stimulus perception

The agent perceives cues in isolation and process them independently, updating their predictive value in accord with Equation 1. However, when a compound of cues is presented, the agent perceives visual features of the compound which are absent when the cues are presented in isolation. For example, as long as cues are presented in close proximity during compound trials (i.e., either overlapping or close to one another), it is possible that the subject will sample visual information in some areas of the display that are unique to compound trials, such as line intersections, corners, etc., which are not available when stimuli are presented in isolation. This makes the compound trial different to a single cue trial by the addition of those features in the configuration. We assume that the agent treats those added features the same as any other cue ^1^.

### Sampling process

Finally, we assume that the probability of stimulus *i* being processed in any given trial is given by a *softmax* function incorporating salience and predictive value of each stimulus (Sanderson and Bannerman, 2011; Sutton and Barto, 1998):

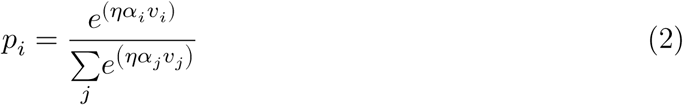

where *η* is a decisiveness or *temperature* parameter that determines the extent to which the agent is biased to sample cues with low salience or predictive value (*j* = [1, 2, …, *i*, …*k*]) ^2^. Finally, in each trial the agent samples from the stimulus array *S* = [*S*_1_, *S*_2_, …, *S*_*k*_] for a single stimulus to process. We assume that the sampled cue *S*_*i*_ follows a categorical distribution:

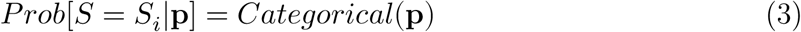

where **p** = [*p*_1_, *p*_2_, …, *p*_*k*_] is the vector of probabilities given by Equation 2 and *k* is the number of stimuli presented in a given trial during training and testing.

To illustrate how this model operates, see the schematic representation of Figure 6. In this example, we assume that a compound AB is presented. In this case, the configuration produces an additional unique cue X which is perceived by the agent for that particular combination of cues. Once the agent samples cue A (shown in red), its value is updated according to the difference between the current value or prediction (*υ*_*i*_) and the outcome observed (*λ*) in that trial. For the following simulations, we assume that the salience of the unique cues is equal to the salience of the target cues; that is, *α*_target_ = *α*_nonTarget_, and that the order of presentation of different trial types is random.

**Figure 6.**
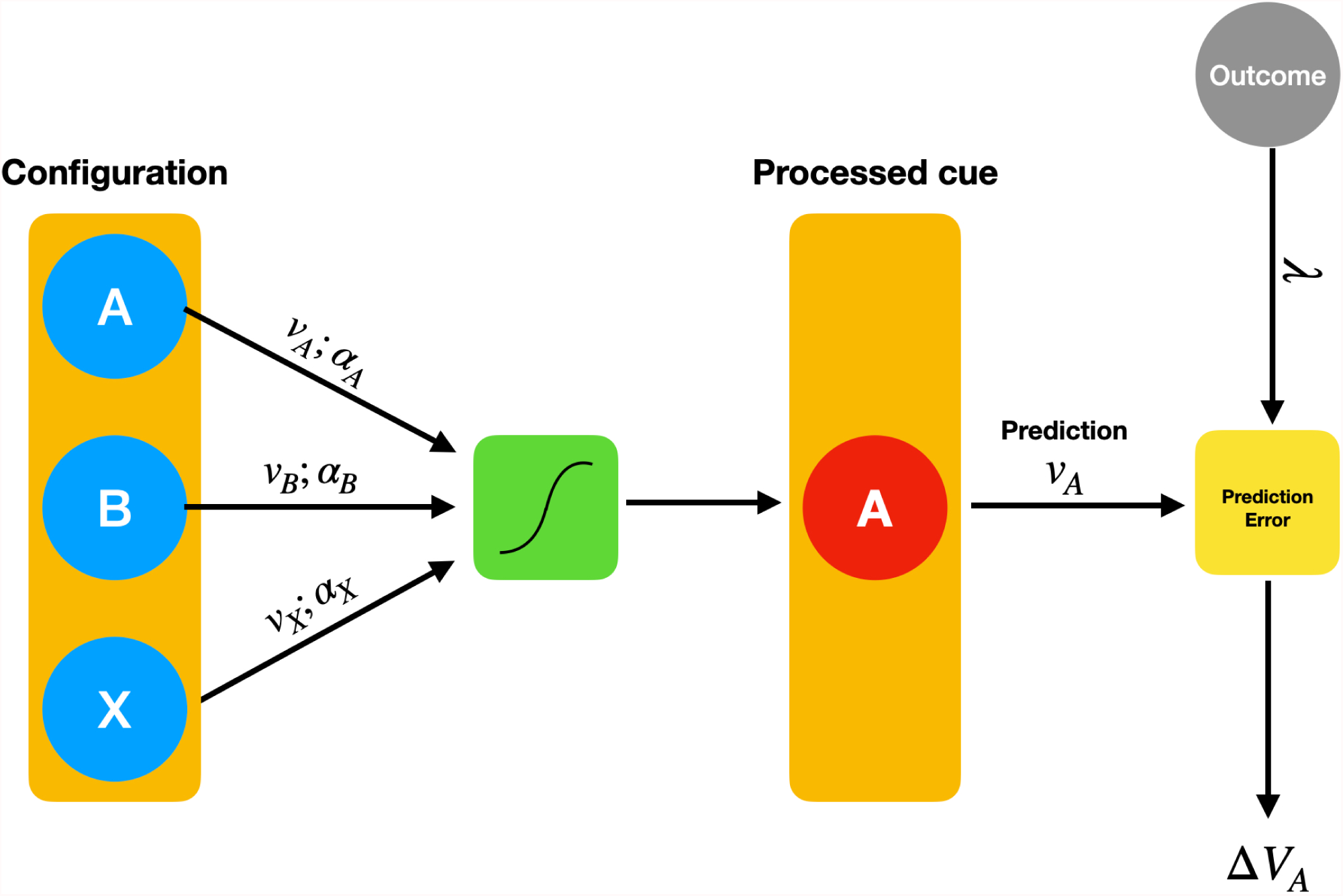
A sub-sampling model. On each trial where the agent is presented with a compound of cues that predict an outcome, the agent searches and samples one of the cues according to their current value (*υ*) and salience (*α*). In this example, the compound AB is presented and an additional cue X is perceived for the compound. For illustrative purposes, we assume that cue A has been sampled in this trial (colored in red). Once the agent samples the cue, it updates its value according to a prediction error rule. Only the value of the sampled cue (A, in this example) is updated for the following trial.

To allow our agent to sample cues with low predictive value we set the temperature parameter *η* to 30 and the initial predictive value of each cue, both target and unique cues, *υ*_*i*_, *i* ∈ [1, …, *k*], to 0.05. Unless otherwise noted, the value of *λ* was set to 1 for reinforced trials and to zero for non-reinforced trials. Finally, we assume that an additional cue for each one of the possible combinations of cues is added when a compound is presented ^3^. We ran 80 simulations for each experimental design. The values shown in the figures are the average values across all simulations.

In addition, note that the model presented here is an attempt to capture only the *serial processing with satisfaction of search* to explain results from groups *extra* and *intra* in our experiments. The parallel processing strategy assumed to happen in group extra2 is not meant to be captured by the model as it is presented here. We assume that traditional elemental models, such as the Rescorla-Wagner model (Rescorla and Wagner, 1972), are applicable under conditions that foster parallel processing (i.e., clearly identifiable component cues). In other words, we assume a model in which only elemental processing of cues occurs, but in which full sampling (i.e., parallel search) or sub-sampling (i.e., serial search with satisfaction of search) of cues makes behavior approximate the predictions of elemental and configural theories, respectively, but we do not provide in this model a decision rule by which the perceptual system deploys full or sub-sampling of cues.

## Results

### Summation

We start by replicating the conditions of a simple summation design in pigeon autoshaping. To this end, we assume for simplicity that both components predict the same outcome value and have equal saliencies (*α*_*A*_ = *α*_*B*_ = .4). The results of this simulation are shown in the right panel of Figure 7a. As shown in the figure, the sub-sampling model correctly predicts the failure to find a summation effect in these experiments. Under the sub-sampling model, the system randomly samples the unique cue that is only perceived in the testing phase (ABX, where X is the unique cue), which tends to bring down responding to AB (ABX) compared to the elements A and B. The left hand panel of Figure 7a shows a similar design reported by Rescorla and Coldwell (1995) using pigeon autoshaping, where the authors failed to find a summation effect.

**Figure 7.**
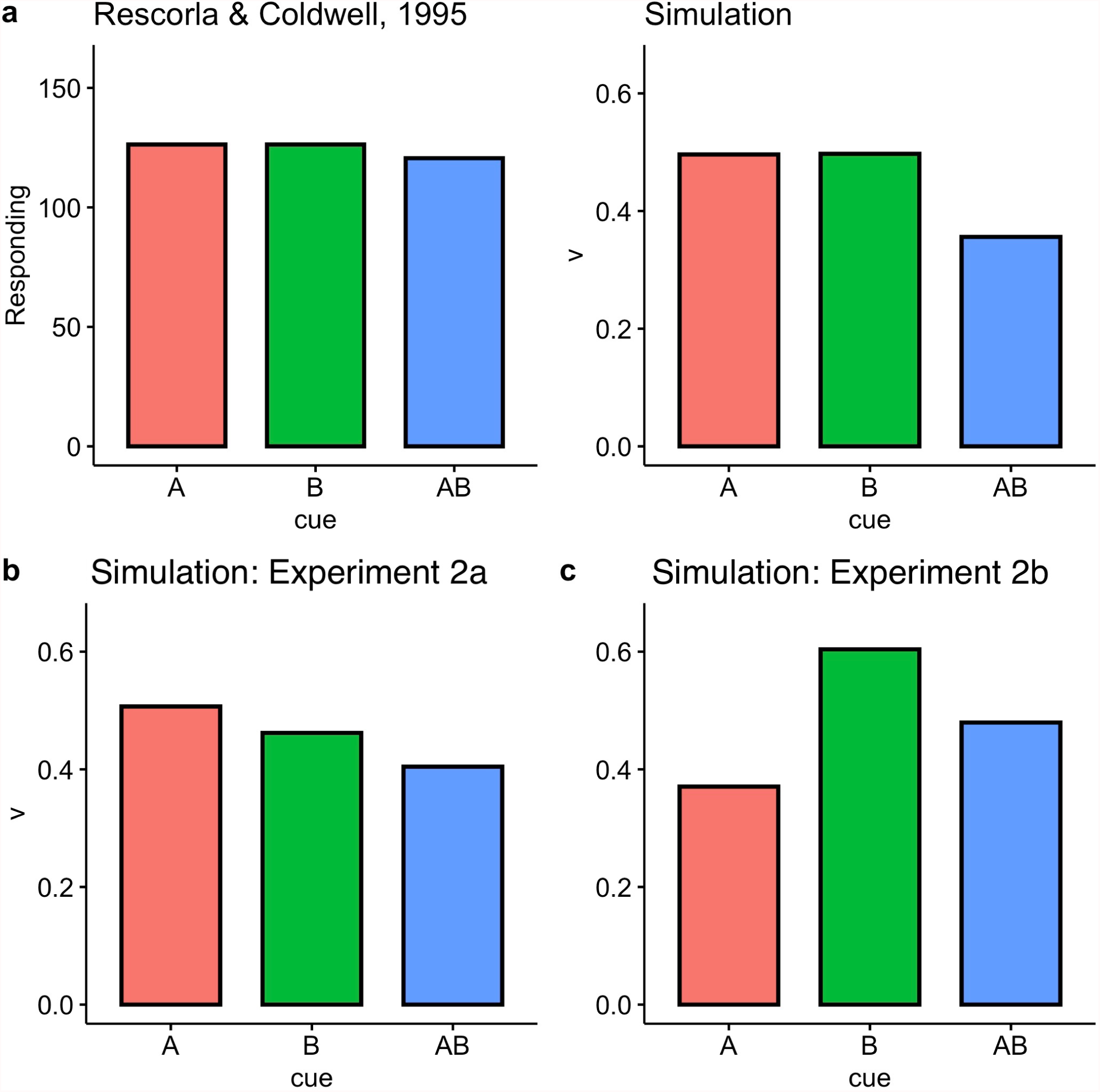
(a) Simulations of the sub-sampling model for different summation designs. The left panel shows the results obtained by Rescorla and Coldwell (1995) in pigeon autoshaping. The right panel shows the simulations of the model assuming that the saliencies of A and B are equal. (b) Simulations of a sub-sampling model for a summation experiment where the salience of B is assumed to be higher than that of A, but B predicts a lower outcome value. (c) Simulations of a sub-sampling model for a summation experiment where the salience of B was higher A but the outcome value predicted by B is higher than the value predicted by A.

In a second set of simulations, we investigated the predictions of a sub-sampling approach when cue B predicts a lower outcome value than A, trying to match the conditions of Experiment 2a. To account for the fact that B seemed to be more salient and drive responding in that experiment, we set the value of *α*_*A*_ to .4 and the value of *α*_*B*_ to .5. A value of .4 was also set to the unique cue X (*α*_*X*_ = .4). To account for different outcome values predicted by each cue, we set *λ*_*A*_ = 1 > *λ*_*B*_ = .95. The model correctly predicts the pattern of results of Experiment 2a, in that responding to A, the most salient cue, is higher than to B (see Figure 7b). The model also replicates our finding that responding to AB would be closer to B than to A. Lastly, we tried to match the conditions of Experiment 2b by reversing the roles of A and B, so that B predicts a higher outcome value than A (*λ*_*A*_ = .95, *λ*_*B*_ = 1). Again, the sub-sampling model correctly captures the pattern of behavioral results in this experiment, anticipating that responding to AB should be closer to the outcome predicted by B, and higher than in Experiment 2a (see Figure 7c).

### Differential summation

In another experiment in pigeon auto-shaping, Pearce and colleagues (1997) found that responding at test for the compound of three cues, ABC, was weaker when the three cues were separately paired with a reinforcer (A+, B+, C+) than when the cues were paired with the same reinforcer, but in compounds (AB+, AC+, BC+) (see Figure 8, left panel). Pearce’s configural model can readily account for these results under the assumption that the similarity between the compounds comprising two cues (AB, AC, BC) and the tested compound ABC is higher than that between each A, B and C and the compound ABC.

**Figure 8.**
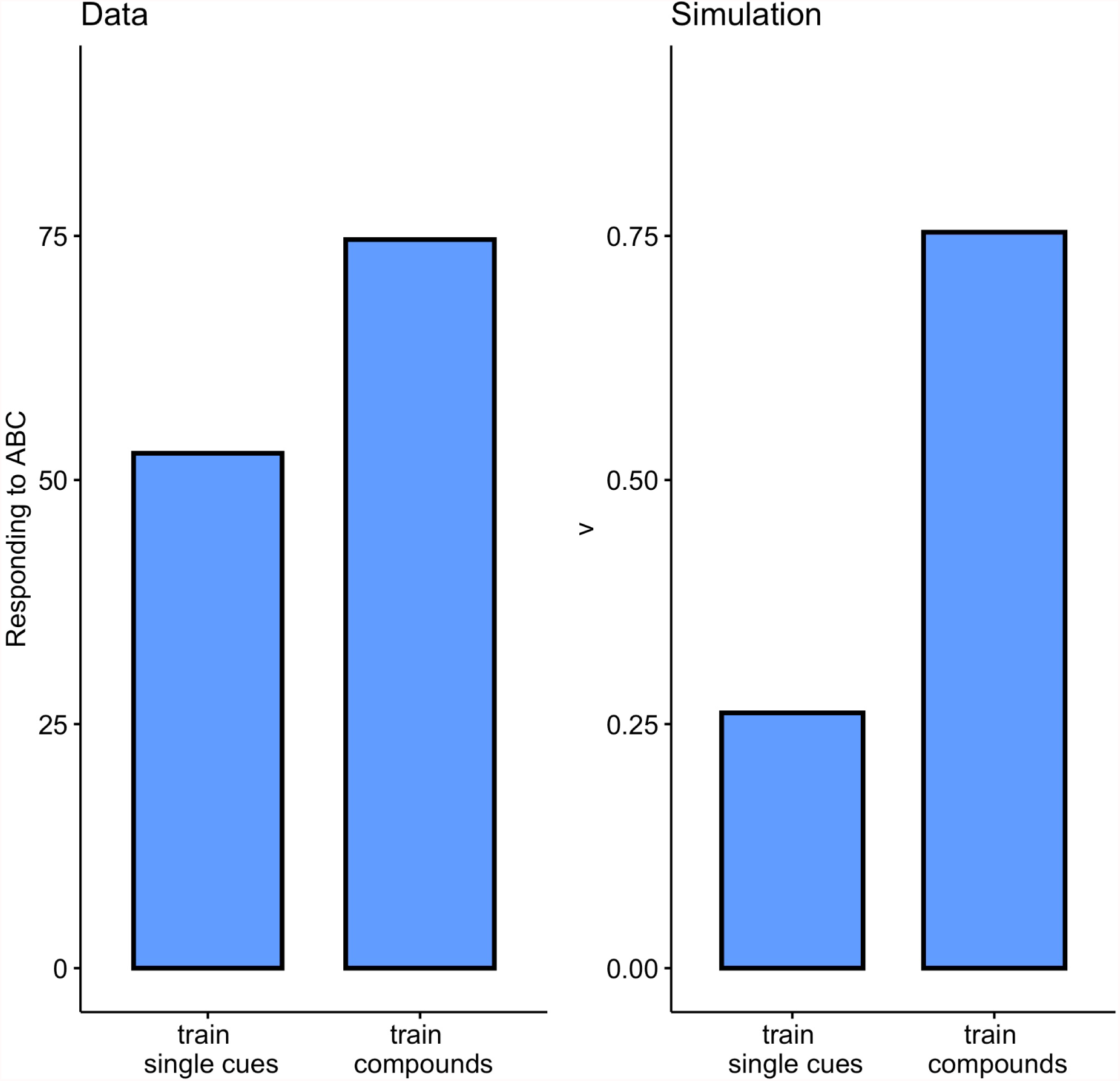
Simulations of a sub-sampling model for a differential summation design. Left panel. Results obtained by Pearce and colleagues (1997) in pigeon autoshaping. The bars show the responding to the compound ABC after training with the single cues A, B and C or after training with the compounds AB, BC, and AC. Right panel. Simulations of the sub-sampling model for the same design.

Figure 8 (right panel) depicts the simulations of this design in the sub-sampling model, which also correctly captures these data. In the case of single-cue training, at the end of training ABC is composed of four unique cues with very low predictive values (.05 each, by assumption), whereas the same ABC compound is composed of the same four unique cues, some of which have been experienced by the agent during training and have therefore acquired higher predictive values than in the case of single-cue training. During the presentation of ABC at test, the agent sometimes sample these higher-valued unique cues. As a consequence, the agent tends to respond more to the compound ABC after compound training than after single-cue training, in agreement with the results observed by Pearce et al. (1997). In other words, our model proposes that it is not the similarity between two-stimuli compounds like AB and the test compound ABC what increases summation in this design, but rather the availability of additional cues perceived by the subject which can be sampled during both training and testing.

### Reversing a conditioned inhibitor

Pearce and Wilson ran an experiment which included a feature-negative design of the form A+, AB-in a first phase, and reinforced presentations of B alone in a second phase (B+; completing a negative patterning design across phases). In the learning literature, the first phase turns cue B to what is referred to as a *conditioned inhibitor*, meaning that B signals the absence of an otherwise present reinforcer. In contrast to an elemental theory which would predict B to recover its predictive value so that responding to AB should be higher than A or B alone at the end of the second phase, Pearce and Wilson observed instead lower responding to AB than A or B alone (see Figure 9, left panel). This result is also anticipated by a sub-sampling approach (see Figure 9, right panel). During the first stage, A acquires more value than both B and the unique cue represented for AB, call it X. During the second stage, B acquires value independently of the sub-sampling process. During the final test with AB, however, the agent still samples the unique cue X, whose value has not been modified during the second phase, staying at a low level. Responding to AB is therefore lower than to either A or B.

**Figure 9.**
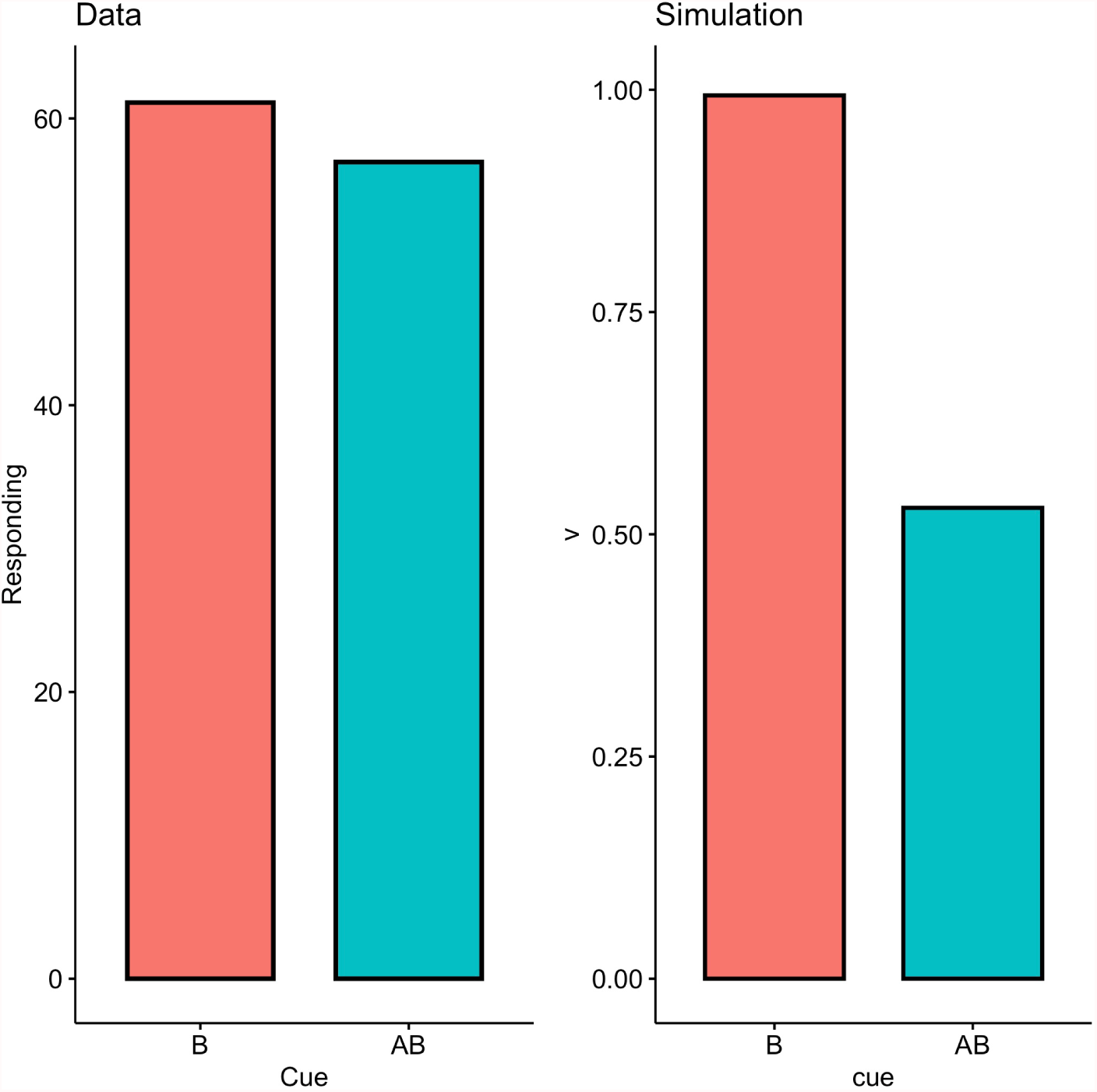
Simulations of a sub-sampling model for Experiment 3 of Pearce and Wilson (1991). The experiment comprises a first phase where A is reinforced and the compound AB is not (A+, AB-). In the second phase B is reinforced in isolation. The left panel shows the original data obtained by these authors. Simulations of the sub-sampling model are shown in the right panel.

## Discussion

This paper reports empirical evidence of similarity affecting summation in humans. In Experiment 1, when stimulus properties were designed to make it difficult for subjects to parse them and deploy counting strategies, the difference in similarity between target and non-target cues within a compound was reflected in different ratings for the compound ABX/ABY during the test. In addition, although previous studies had demonstrated that summation could disappear when participants assumptions are affected by prompting rational rules concerning the independence of cues in a compound (Pérez *et al*., 2018), we observed null summation in the absence of such manipulations. We replicated this absence of summation in three different experiments.

To our knowledge, only one previous paper by Lachnit (1988) obtained evidence for similarity affecting summation. Consistent with the animal evidence employing unimodal versus multimodal stimuli, Lachnit tested compounds of visual stimuli that were “separable”, varying in size and orientation, and observed a level of summation similar to that predicted by elemental theory. By contrast, when he used target stimuli considered to be “integral”, varying in saturation and brightness, the summation effect became weaker and closer to the predictions of configural theory. Although the result appears to be similar to ours, there are important differences between the designs. In Lachnit (1988), the stimulus dimensions belonged to a single object whereas our stimuli were presented in compound during training and the similarity was varied between target and non-target cues. Our approach is therefore not directly applicable to the type of stimuli employed by Lachnit (1988). However, both papers illustrate the need to incorporate what is known of perceptual mechanisms affecting the input to the associative learning machinery.

A notable result in Experiment 1 was that the driver of the summation effect was brought about by the similarity between the target cues A and B and non-target cues X and Y, which contrasts to the notion of similarity proposed by learning models, which assume that the driver should be the similarity between the target cues. Figure 10 shows the expectations for Experiment 1 based on the assumption of an effect of similarity on summation in associative models. If similarity between cues A and B is the critical variable driving summation, we should have observed higher summation in groups *extra* and *extra2*, where A and B are dissimilar, than in group *intra* (Figure 10; left panel). If, in addition, similarity between the targets A and B and non-target cues X and Y influences summation, we should have obtained higher summation in group *extra2*, where those cues are dissimilar, than in group *extra*, and higher summation in group *extra* than in group *intra* (Figure 10; right panel.)

**Figure 10.**
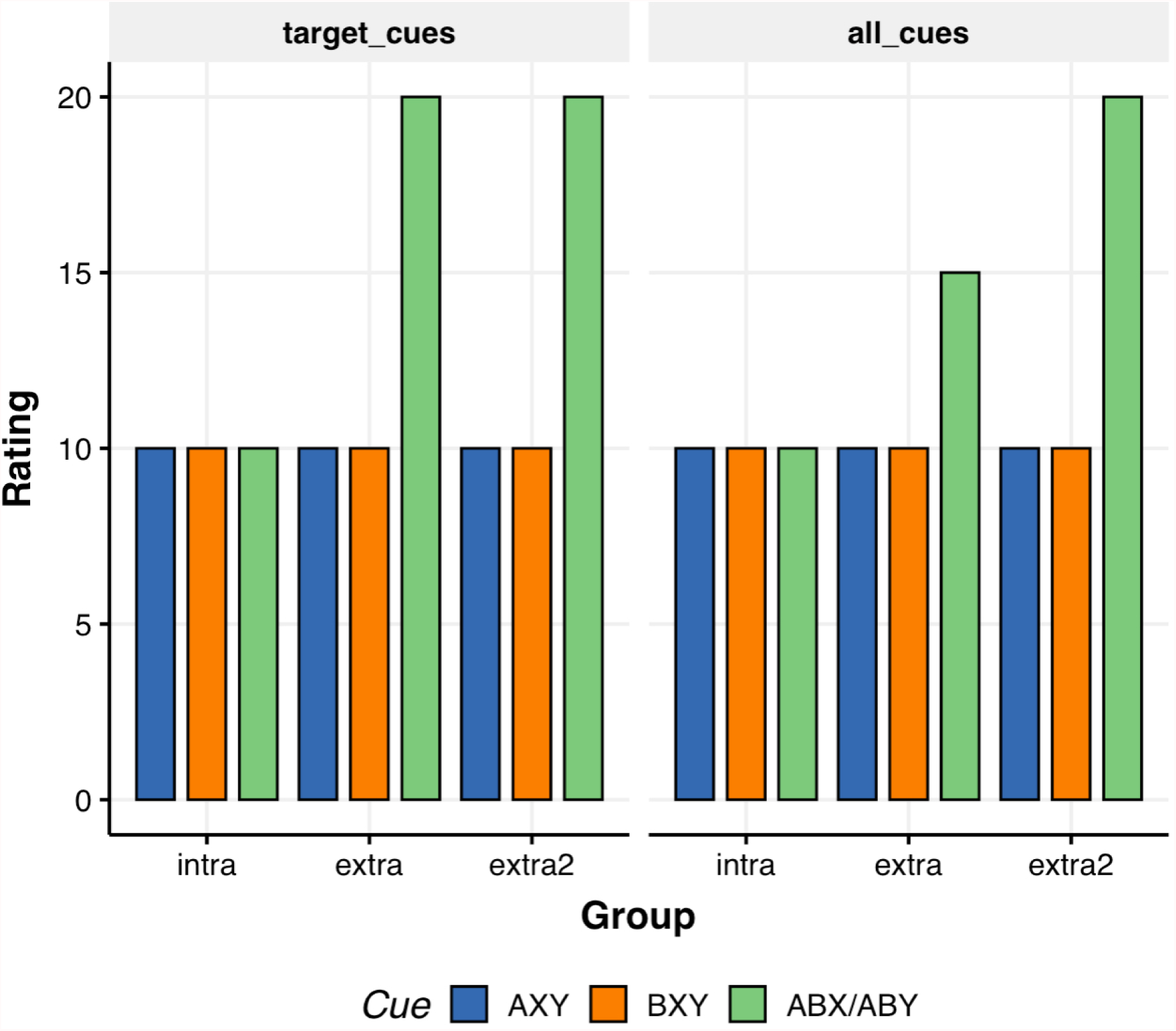
Expectations for Experiment 1, based on the prediction from associative learning theory of an effect of similarity on summation. **Left panel.** Expected results when similarity of only the target cues A and B influences summation. **Right panel**. Expected results when similarity of all cues (i.e., non-target cues as well as target cues) in-fluences summation.

To confirm these predictions through a more formal associative analysis, we performed simulations with one of the most flexible of associative models, the Replaced Elements Model (REM) proposed by Wagner and his colleagues. In short, REM assumes that the presentation of a given cue activates two types of elements: Context-independent and context-dependent elements. The context independent elements are activated whenever the target cue is presented independent on whether other cues are present or absent. In turn, context independent elements, apart from the cue they represent, are activated only when other specific cues are present or absent. In the case of our experiments, for instance, when ABX is presented at test, context-independent elements *a*_*i*_, *b*_*i*_ and *x*_*i*_, and context-dependent elements *a*_*x*_ and *b*_*x*_ contribute to the prediction. At the same time, some predictive value is also lost when ABX is presented due to the replacement of several excitatory elements by associatively neutral elements. The overall result, of course, will depend on the assumptions made about the replacement between components, but the predominant result in REM is negative summation. Indeed, we could not reproduce the results of Experiment 1 even when testing a wide range of values in its parametric space (see Brandon *et al*., 2000; Vogel *et al*., 2017; Wagner, 2008 for theoretical details and Appendix for results of those simulations).

In Experiments 2a and 2b we obtained evidence for the sub-sampling hypothesis by showing that the majority of participants rated a compound ABX/ABY equal to the value of one of the cues in the compound. Again, associative models cannot explain these results as they predict lower responding to ABX/ABY in groups *intra2* than in group *intra*—in the former group the asympotic values for AXY and BXY are 8 and 10, respectively, whereas these values are both 10 for group *intra*. Moreover, when we swapped the outcome values predicted by AXY and BXY between Experiments 2a and 2b, participants followed the value of one of the stimuli, BXY, presumably due to it being more salient.

Another interpretation for these results can be made by examining further potential visual mechanisms. One possibility is considering a perceptual grouping hypothesis^4^, depicted in the left panel of Figure 11. As already described, the cues in our design varied in shape only and were all composed of eight circles connected by lines. Given that the three different shapes in a compound were connected with one another via the central circle, perceptual grouping of cues of the same color may result in them being perceived and encoded as a single configuration (Palmer, 1999). In Figure 11, one can see that for group *intra* this would result in encoding of a single three-cue group during training, and encoding of a novel but similar three-cue group during testing. For this group, therefore, one would expect a rating for the testing compound ≤ 10, with the actual value depending on how much learning about the training group one assumes generalizes to the testing group.

**Figure 11.**
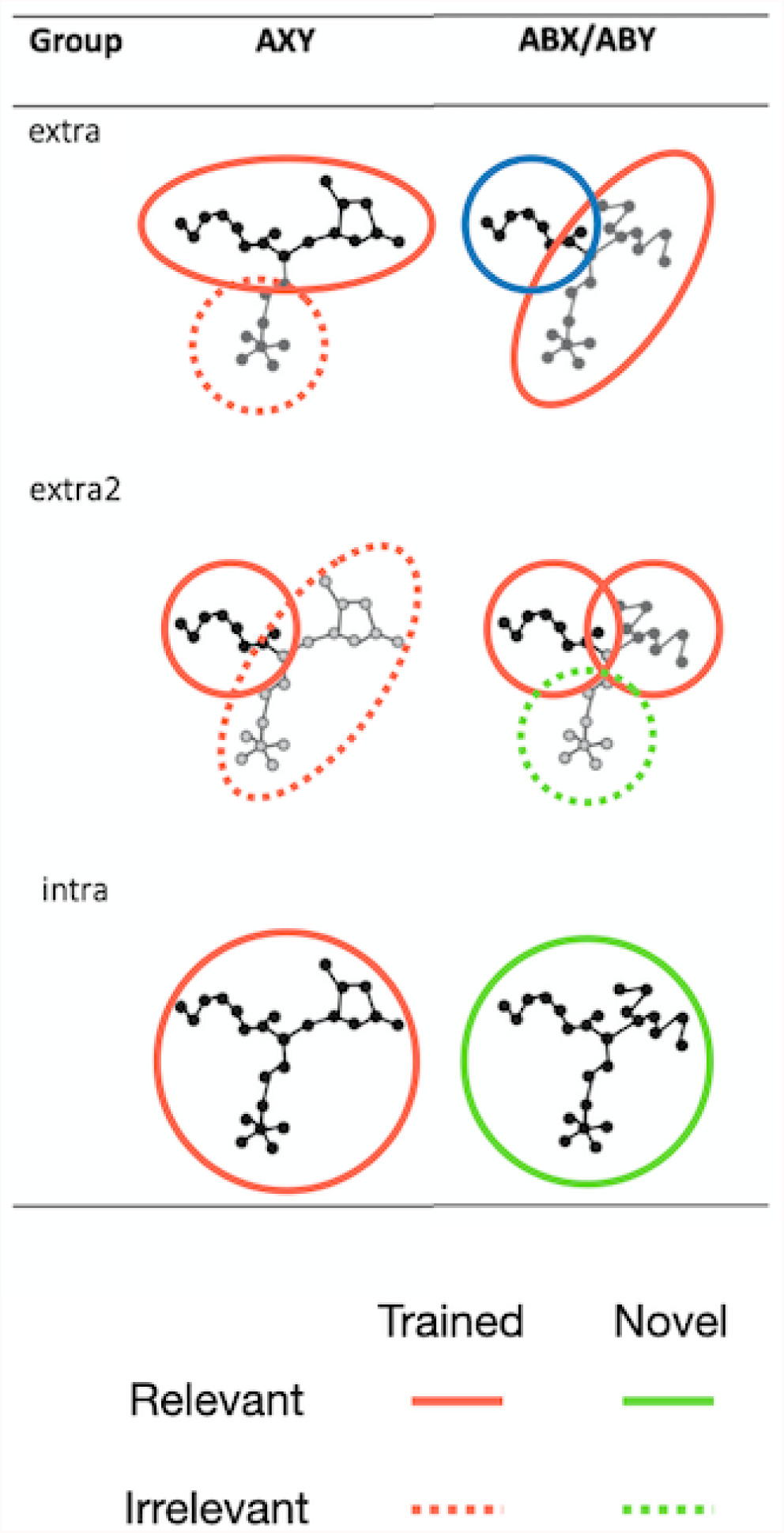
Perceptual Grouping hypothesis for Experiment 1 (see main text for details).

The predictions are more complex for groups *extra* and *extra2*. In group *extra*, during training people should parse the compound into two groups: one composed of a target cue and a non-target cue, both being relevant for outcome prediction (solid red ellipse in Figure 11), and one composed of a single non-target cue, being irrelevant for outcome prediction (dotted red circle in Figure 11). The testing compound would be parsed again into two groups: one being a trained relevant group of two cues, associated with 10 degrees of allergy (solid red ellipse in Figure 11), and one being a novel group composed of a single target cue (green circle in Figure 11). For this group, one would expect a rating for the testing compound > 10, with the actual value depending on how much learning about the two-cue training group one assumes generalizes to the single-cue testing group. Finally, in group *extra2*, during training people should parse the compound into two groups: one composed of the target cue, being relevant for outcome prediction (solid red circle in Figure 11), and one composed of the two non-target cues, being irrelevant for outcome prediction (dotted red ellipse in Figure 11). The testing compound would be parsed into three groups of single cues, two of them being trained groups associated with ten degrees of allergy. For this group, one would expect a rating for the testing compound ≈ 20. In sum, the predictions of this perceptual grouping hypothesis are that ratings to the ABX/ABY testing compounds should be: *extra*2 ≈ 20 > *extra* > *intra* ≤ 10. Thus, the perceptual grouping hypothesis is more than superficially similar to configural theory, as it makes the same rank predictions as all versions of REM implementing some configural processing (see Appendix).

In the last section of the paper, we presented a computational implementation of the sub-sampling hypothesis. Through simulation we showed how this model is able to capture not only the data of groups *intra* and *extra*, but also a wider range of previous phenomena that are usually attributed to configural processing of stimuli in the animal literature. In contrast to these models, however, we obtained these results by taking a view that relies on objective properties of stimuli—some of which are absent when stimuli are presented in isolation—rather than on internal representations, as it is usually assumed in associative learning models, which makes this approach more parsimonious than current theories based on internal stimulus representations.

The sub-sampling model outlined in this paper does not contain a mechanism for sampling multiple cues in a compound and is therefore not meant to capture summation data. We have assumed that traditional elemental models, such as the Rescorla-Wagner model (Rescorla and Wagner, 1972), can explain behavior under conditions that foster parallel processing during visual search (i.e., clearly identifiable component cues). A simple extension could be formulated to formalize a mechanism that chooses between sub-sampling and full sampling strategies. As a first approximation, the mechanism could be implemented in an all-or-none fashion. For example, the agent could arbitrate between strategies following a rule such as

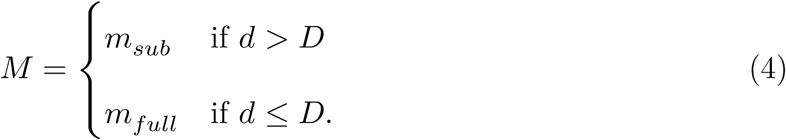

where *d* is a parameter that determines how difficult is to parse components—which should be a direct function of the similarity between them—and *D* is a threshold that varies across individuals and which will determine whether sub-sampling or full sampling is deployed.

Regardless of the possible extensions of a sub-sampling approach, our work provides empirical and computational evidence suggesting that under conditions in which components in a compound are difficult to parse and identify, participants may engage in an inefficient search process by which they only sample a subset of the compound. This sub-sampling approach can explain the empirical evidence presented in this paper better than current models of learning, and a simple formalization of its principles can also capture previous data from the animal literature usually interpreted as supporting configural processing. We believe this interpretation is testable, important, and should be considered as a plausible alternative hypothesis in the literature on learning and generalization.

## Materials and Methods

Participants were tested in desktop Windows (c) computers running Psychopy (version 1.75 for Experiment 1; version 1.82.4 for Experiments 2 and 3; Peirce, 2007). Responses were recorded from standard PC keyboards.

### Statistical analysis

Statistical analyses were performed using RStudio and SPSS 27.0. For all the pre-planned comparisons we calculated a Welsh *t*-test and included Cohen’s D, along with a 95% confidence interval on this estimate, as a measure of effect size. When reporting interactions between factors, we computed 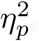. Following Steiger (2004), we report a 90% confidence interval on this estimate. The reliability of the results was contrasted against the usual criterion of *α* = .05. When a null effect was expected, a Bayes Factor in favor of the null (*BF*_01_) is reported.

## Experiment 1

### Participants

Eighty-six undergraduate students from Florida International University participated in Experiment 1. Participants did not have previous experience with the experimental procedure and were tested simultaneously and in the same room. The Institutional Review Board of Florida International University approved all the studies in this paper (IRB-15-0460). Written informed consent was obtained from all participants. They were given course credit for their participation.

Participants were randomly assigned to one of three groups: *intra* (*n*_*intra*_ = 27), *extra* (*n*_*extra*_ = 28) and *extra2* (*n*_*extra*2_ = 31). In all of the studies in this paper, we determined our exclusion criteria before data collection by following the criteria employed in Pérez *et al*., 2018. Participants that did not give on average a rating between 7 and 13 points of allergy to cues that predicted allergy and between 0 and 3 points of allergy to cues that did not predict allergy, were left out from the analysis. The final number of participants in Experiment 1 was *n*_*intra*_ = 18, *n*_*extra*_ = 13, *n*_*extra*2_ = 27.

### Procedure

Before the training phase, participants were presented with the following instructions:

> *In this experiment, we will ask you to imagine that you are an allergist; that is, a medical specialist whose work is to discover the causes of allergic reactions in people. A new patient, Mr. X, visits you asking for your help. Mr. X has allergic reactions to some drugs, but not to others. In an attempt to discover which drugs cause allergic reactions to Mr. X, you apply the drug on Mr. X’s skin and observe the magnitude of the allergic reaction. On each trial, the computer will show you a shape representing the drug that has been applied to Mr. X’s skin. You will be asked to predict the magnitude of the allergic reaction that Mr. X will have to the drug. Enter your prediction using the rating scale that will be shown at the center of the screen. This scale ranges from 0 to 35 points, where 0 points means that the drug produces no allergic reaction and 35 points means that that the allergic reaction is of maximum intensity. You can adjust your prediction as many times as you want and take as long as you like to enter the prediction. Once you feel satisfied with your prediction, you must confirm it by either pressing the “Enter” key in your keyboard or clicking on the grey button below the rating scale. You will receive feedback about the actual magnitude of the allergic reaction. At first, you will need to guess the correct answer, but your predictions should get more accurate as the experiment progresses*.

Participants were presented with the following instructions before the test phase:

> *Now you will have to make a final evaluation of the drugs that produce allergic reactions in Mr. X. The computer will show you single drugs or combinations of drugs and, as before, you must enter your prediction using a scale that ranges from 0 to 35 points, where 0 points means that the drug produces no allergic reaction and 35 points means that the drug produces an allergic reaction of maximum intensity. Use intermediate values to indicate different degrees of allergic reaction between no allergic reaction and maximum allergic reaction. To confirm your choice, press “Enter” in your keyboard or click the gray button that will be shown below the rating scale. You can change your decision as many times as you want before confirming it*.

Groups differed in the similarity between cues in the display (see Figure 2). Each stimulus was created from three different cues that “branched out” from a central point. Among these branches, only one of them represented the target cue associated with either allergy or no allergy during training (A, B, C or D). The other two branches were non-target cues that could not predict the presence or absence of allergy (X and Y). During the test, the compound AB was comprised by two target branches together with an additional non-target cue (ABX or ABY). In group *intra*, all these “branches” were of the same color (black), but differed in shape. In group *extra*, A and B differed in color (grey and black), but they shared color with the non-target cues (X and Y, one grey and one black). In group *extra2*, the target cues were the same as in group *extra*, but now the non-target cues had a distinctive color as well. In all groups, A and B, which predicted allergy, shared color with cues C and D, which predicted no allergy. Thus, all participants, regardless of group, had to attend to shape; color was irrelevant to solve the discrimination.

## Experiment 2a

### Participants

75 undergraduate students from Florida International University were randomly assigned to one of two groups (*n*_*intra*_ = 40, *n*_*intra*2_ = 35) and were compensated with course credit for their participation. The exclusion criteria were the same as those of Experiment 1, except that the admittance interval for the mean rating to A and B in group intra2 was set to [5-11]. The final number of participants per group was *n*_*intra*_ = 39, *n*_*intra*2_ = 33.

### Procedure

The procedure was the same as described for group *intra* of Experiment 1, with only one exception: In group *intra2*, stimulus BXY was associated with 8 points of allergy during training (see Figure 5a).

## Experiment 2b

### Participants

80 undergraduate students from Florida International University were randomly assigned to one of two groups (*n*_*intra*_ = 42, *n*_*intra*2_ = 38) and were tested under the same conditions of Experiment 2a. The final number of participants per group was *n*_*intra*_ = 24, *n*_*intra*2_ = 33.

### Procedure

The procedure was the same as in Experiment 2, except that the outcome values for A and B were interchanged in group *intra2*. In this experiment, the outcome assigned to BXY was 10 while a value of 8 was assigned to AXY.

## Acknowledgements

We thank Jaron Colas, Tomislav Zbozinek and Tony Dickinson for their valuable comments on a previous version of this manuscript. S.D.G.

## Open Practices Statement

The data and materials for all experiments are available at https://osf.io/xqnpk/

## Footnotes

1 A similar idea is proposed in Wagner and Rescorla (1972) and REM Brandon *et al*., 2000; Wagner, 2008). However, in contrast to these models, we do not rely on internal representations but simply assume that objective properties of stimuli are perceived when compounds of cues are presented in close spatial proximity.

2 Formally, the model should include the absolute value of *υ*, since cues with high inhibitory strength (negative *υ*) should command more attention than other cues in a given array (Parkhurst *et al*., 2002). However, such implementation is only relevant in a model that allows for negative associative values for cues, which is not the focus of our model.

3 For example, if the compound ABC is presented, we assume that there is an additional unique cue represented for each possible pair—AB, AC, BC—and the compound of three cues—ABC. The compound ABC is thus represented as ABCXYZV, where X, Y, Z, and V are the additional cues which are perceived by the agent.

5 We thank Harald Lachnit for mentioning this possibility.

